# Comparative ubiquitinomics of human skin reveals insulin receptor ubiquitination as a regulator of collagen secretion

**DOI:** 10.64898/2025.12.03.691868

**Authors:** Joseph Inns, Andrew Michael Frey, Ishier Raote, Mads Gyrd-Hansen, Matthias Trost, Neil Rajan

## Abstract

Ubiquitination is central to skin homeostasis and disease, however the cutaneous ubiquitinome remains poorly characterised. We investigated spatial patterns of ubiquitination in human skin, discovering specific ubiquitin linkages located within the hair follicle and epidermis. To create a comprehensive physiological profile of the ubiquitinated proteins within the skin, we profiled healthy skin and CYLD cutaneous syndrome (CCS) skin tumours through di-glycine remnant immunopurification proteomics, enabling identification of 1605 ubiquitin sites across 731 proteins. The healthy cutaneous ubiquitinome comprised substrate classes including keratins, S100 proteins and histones. Predictive upstream enzyme analysis ranked the most frequent E3 ligase as NEDD4, and the most frequent deubiquitinases included USP7 and CYLD. Analysis of the CCS tumor ubiquitinome, characterized by catalytically inactivated *CYLD* somatic mutations and prominent extracellular matrix (ECM) secretion, identified 251 differentially ubiquitinated sites compared to healthy skin. The CCS ubiquitinome was enriched for cell division and differentiation proteins, including predicted CYLD substrate insulin receptor (INSR). Secretome analysis of INSR knockdown CCS primary cells revealed reduced secretion of basement membrane proteins, particularly COL7A1. By defining how ubiquitination modulates pathophysiological ECM secretory output, our study highlights the ubiquitin post-translational system as a broader determinant of tissue architecture.

**Graphical abstract:** 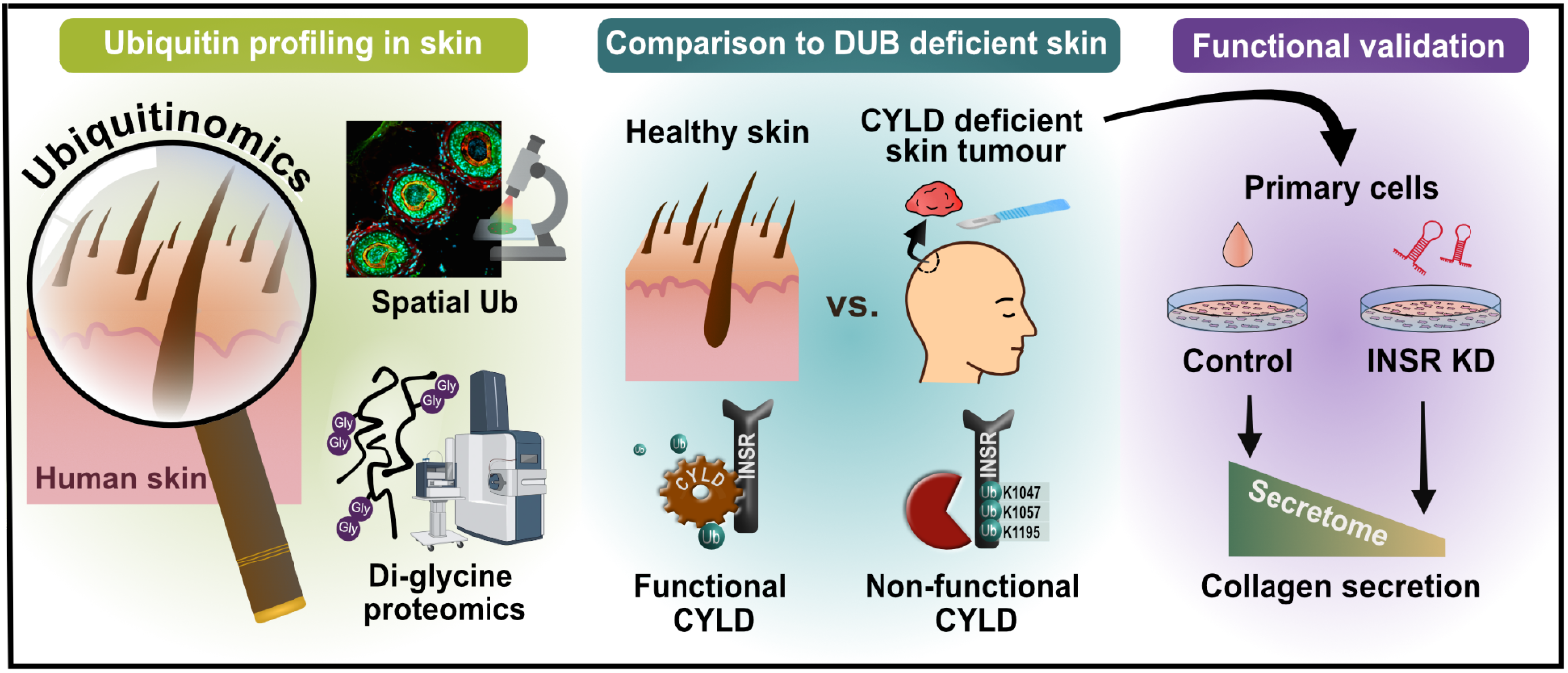

## Introduction

Protein turnover, localisation, and conformation are regulated by post-translational modifications (PTMs), of which ubiquitination is one of the most frequent and functionally diverse.^1^ Ubiquitin (Ub), a highly conserved 76 amino acid protein, modifies target proteins by conjugation to lysine residues. Ubiquitination is multifarious: ligated as a single moiety (mono-ubiquitination), or as poly-ubiquitin chains assembled upon Ub’s internal lysine residues (K6, K11, K27, K29, K33, K48, K63) or N-terminal methionine (M1), with each Ub type promoting a distinct cellular outcome^1^. Ub PTMs are hence capable of directing varied biological processes including cell cycle progression, DNA damage response, immune signalling, trafficking, and chromatin remodelling^2–6^.

The importance of Ub ligation and deubiquitination, performed by ubiquitin E3 ligases (E3s) and deubiquitinases (DUBs) respectively, is underscored by phenotypes arising in patients with pathogenic variants (PVs) in these enzymes. PVs in DUBs encoded by *PRKN* are associated with Parkinson’s disease, *BRCA1* with breast cancer predisposition and *USP26* with male infertility^7–9^. In the skin, PVs of E3s *MDM2*, and *KLHL24* can lead to Lessel-Kubisch syndrome and epidermolysis bullosa respectively^10,11^. Additionally, PVs in the DUB, *BAP1* are associated with melanoma development risk^12,13^. In CYLD cutaneous syndrome (CCS), patients develop hair follicle tumours driven by biallelic inactivating somatic mutations of the DUB *CYLD*, resulting in catalytic inactivation^14,15^. Whole genome sequencing of CCS tumours have demonstrated CYLD loss of function as the key driver^15^, making CCS tumours a human “knockout” model and natural perturbation to investigate gene function of this important DUB. Morphologically, CCS tumours are characterised by repeating cylinders of epithelial tumour cells encircled by an abnormal, thickened basement membrane; CCS cells hypersecrete large amounts of extra-cellular matrix (ECM)^16^, including COL7A1^17^ and laminins^18^. While CYLD’s role in negatively regulating NF-κB signaling *via* deubiquitinating substrates including TRAFs, TAK1, IKK gamma^19^ and RIP1^20^ is characterised in several cell line and mouse models, the ubiquitinome of human CYLD inactive tumours remain unexplored^21^.

We performed ubiquitinome profiling of healthy and CCS skin samples to investigate how Ub substrates are altered in a relevant disease context. Liquid chromatography tandem mass spectrometry (LC-MS/MS) of immuno-precipitated di-glycine remnant (K-ε-GG) peptides enabled unbiased, proteome-wide profiling of ubiquitin and ubiquitin-like modifications, where >94% peptides are thought to represent ubiquitin conjugation^22^, an approach recently demonstrated in a proof-of-principle study in a human skin biopsy^23^. By comparing the ubiquitinome of CCS tumours to healthy skin, we found increased INSR ubiquitination in CCS cells. We demonstrate INSR knockdown in CCS primary cells alters the cellular secretome and reduces COL7A1 secretion and other basement membrane proteins. We propose aberrant INSR ubiquitination in CYLD deficient tumours underpins the hypersecretory phenotype of CCS tumour cells and reveals a role for ubiquitination in basement membrane homeostasis and the specification of tumour architecture.

## Results

### Spatial enrichment and localization of ubiquitin within the epidermis and hair follicle

Skin cell fate is determined by Ub PTMs^24,25^, and hence we sought to spatially characterise ubiquitin linkages in human skin. First, we assessed the localisation of total Ub conjugated proteins utilising the FK2 Ub antibody across three different skin regions: the epidermis, hair follicle and eccrine glands. We observed that the FK2 Ub-antibody, which recognises mono-ubiquitination accounting for > 90% of total Ub in most tissues^26^ and some polyubiquitin chain types, localised at the nuclear region of suprabasal keratinocytes in the epidermis (**Figure 1A**), with more intense nuclear staining in the upper epidermis (**Figure 1B**). There was a strong correlation between conjugated Ub and DAPI staining (r^2^ =0.958). Conversely, by using antibodies specific for M1 and K63 Ub chains, we detected these subtypes in cytoplasmic regions of the epidermis (**Figure 1C-D**), suggesting these forms of ubiquitination play roles in processes such as membrane signalling and trafficking rather than roles related to transcription, typically associated with mono-ubiquitination. Ub localisation similarly in the eccrine coil and secretary cells in human skin was also predominantly nuclear (**Figure 1E**). In contrast within hair follicles, both FK2 Ub staining and K63 Ub staining were largely cytoplasmic and prominent in the inner root sheath, where DUBs such as CYLD have been reported to be highly expressed^27^ (**Figure 1F**). Intriguingly, the pattern of K63 Ub was distinct to that detected by the FK2 antibody, with enrichment in the cytoplasm of the cuticle and in the dermal sheath, highlighting K63 Ub enriched substrates within hair follicles^28^.

**Figure 1.**
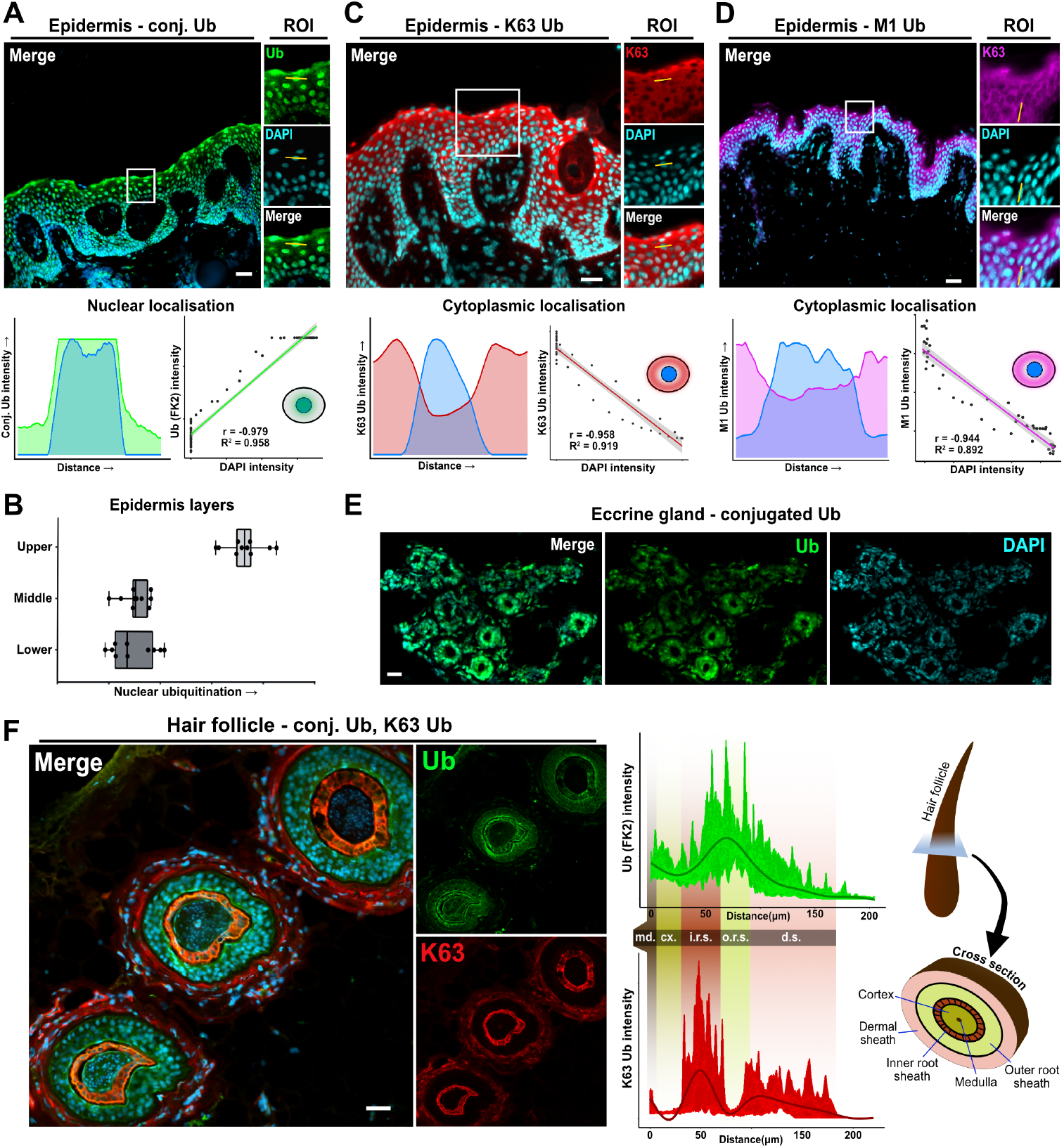
Spatial enrichment and localization of ubiquitin within the epidermis, eccrine gland and hair follicle. **A)** The epidermis is enriched with ubiquitinated proteins with prominent nuclear localisation in keratinocytes. Upper panel - immunofluorescence of human skin sections utilising the FK2 Ub antibody (Green) and DAPI (Blue); Lower left panel - fluorescent image intensity analysis across single cells (yellow line) indicating total Ub and DAPI (n=50 cells); Lower right panel - correlation analysis of total Ub expression and nuclear localisation signal. **B)** Ubiquitination measured by immunofluorescence (FK2 Ub antibody) is increased at the upper epidermis in keratinocytes. Error bars represent standard deviation. **C)** Specific Ub chain types including M1 and **D)** K63 are also detected in the epidermis. Upper panels - immunofluorescence of human skin sections utilising a **C)** K63 Ub antibody (red) and a **D)** M1 Ub antibody, as well as DAPI staining (Blue); Lower left panel - fluorescent image intensity analysis across single cells indicating Ub chain intensity and DAPI (n=50); Lower right panel - correlation analysis of Ub chain intensity and nuclear localisation signal. **F)** Left panel; Hair follicles are sites of ubiquitination with enrichment of K63 Ub chains at the inner root sheath. Right panel; Ub FK2 (green), and K63 Ub (red), immunofluorescence intensity was measured along multiple lines from the centre of the hair follicle outwards. m.d = medulla, cx. = cortex, i.r.s. = inner root sheath, o.r.s = outer root sheath, d.s = dermal sheath. The scale bar is 50 μm.

### The healthy human skin ubiquitinome

To characterise the protein targets of ubiquitination in skin, we extracted protein from human whole skin sections and performed di-glycine (K-ε-GG) immunoprecipitation of the trypsin digested peptides for LC-MS/MS analysis. This workflow (**Figure 2A**) allowed us to analyse the Ub and Ub-like modified proteins in human skin (n=5), where we identified 1465 physiologically relevant K-ε-GG peptides in human skin belonging to 674 proteins. We matched 529 of these proteins to proteins we previously identified in a human skin proteome (n=6) and an additional 145 proteins were only identified after K-ε-GG enrichment (**Supplementary data 1**). We ranked the most prevalently ubiquitinated proteins by their ubiquitination level after normalisation of each protein to its matched skin proteome protein intensity (**Figure 2B, Supplementary data 2**). The highest level of ubiquitination was in SPART, a protein which regulates lipid droplet turnover and epidermal growth factor receptor (EGFR) degradation^29,30^. Cell surface receptors including GPRC5C and DDR1 were amongst the most highly ubiquitinated proteins in skin, as well as ubiquitination related to DUBs such as YOD1.

**Figure 2.**
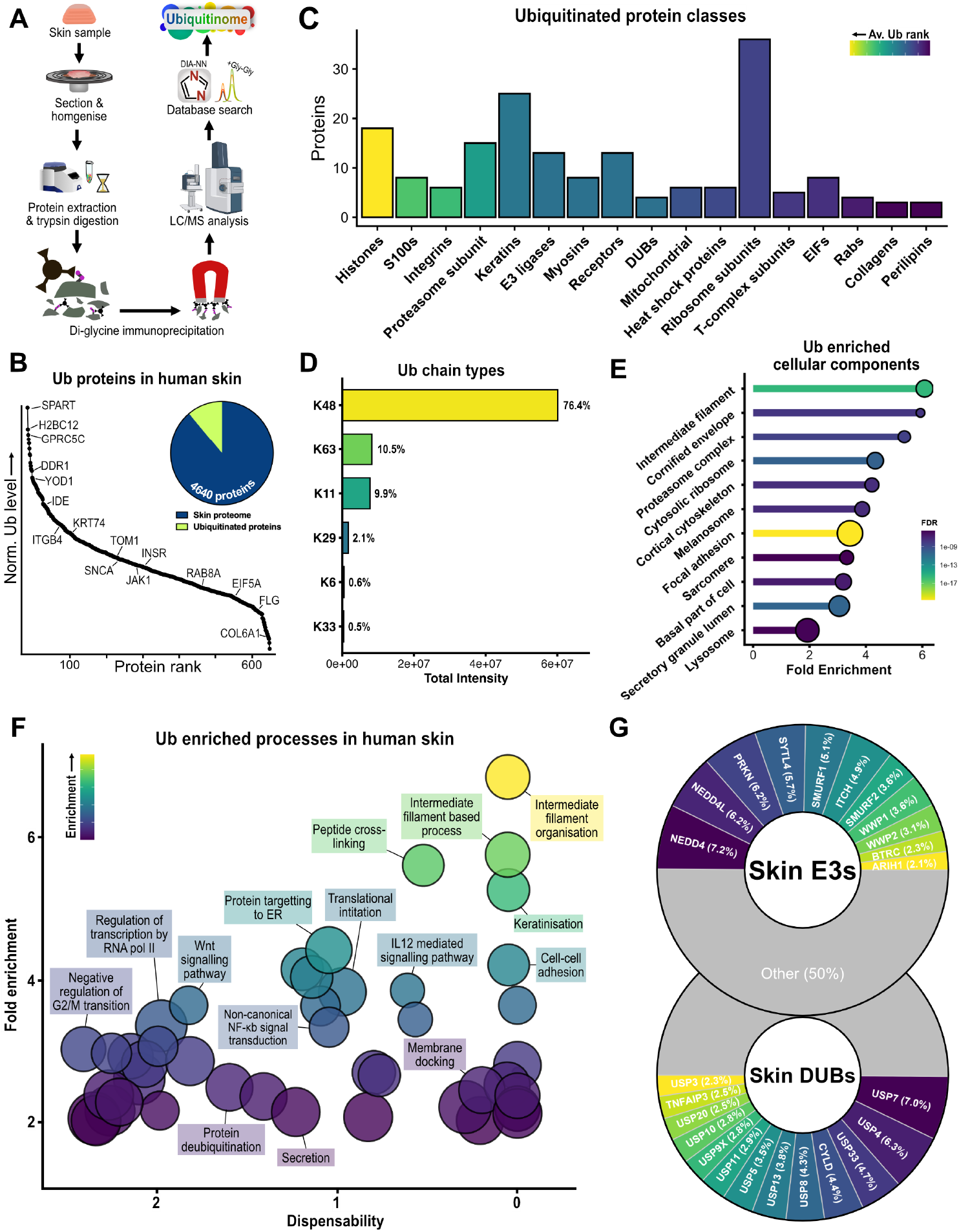
The healthy human skin ubiquitinome reveals widespread substrates and cellular signalling pathways that are regulated by ubiquitin. **A)** The workflow for generating a skin ubiquitinome involved sectioning human skin tissue for protein extraction. Trypsin digested peptides were immunoprecipitated with an anti K-ε-GG antibody and analysed by LC-MS/MS and DIA-NN software. **B)** Skin proteins of K-ε-GG remnant peptides normalised by their matched proteome intensity in rank order. **C)** The protein classes most frequently detected with ubiquitinated sites in skin. Colour indicates increasing ubiquitination calculated by rank. **D)** Prevalence of poly-ubiquitin chain types was assessed by measuring chain type signature ubiquitin peptides. **E)** The ubiquitinated proteins in human skin were enriched for cellular component gene ontology terms. Dot size shows relative number of genes in term. **F)** Enriched Biological Processes of ubiquitinated skin proteins were summarised using REVIGO and plotted to show enrichment, dispensability (similarity to other terms) and GO term size (circle size). **G)** Ubibrowser was utilised to predict ubiquitin E3 ligases and deubiquitinases responsible for ubiquitination/ deubiquitination of detected peptides.

To understand the diversity of ubiquitination in skin we looked at Ub levels across protein classes important to skin function (**Figure 2C**). We observed that some protein classes were frequently ubiquitinated, such as histones, integrins, keratins, and ribosome subunits. H2BC12, whose ubiquitination is associated with active chromatin and transcriptional elongation^31^, was the most highly detected histone. In contrast collagen proteins were less frequently ubiquitinated. Furthermore, we investigated the prevalence of Ub chain types in human skin by measuring the intensity of Ub peptides with K-ε-GG remnant motifs (**Figure 2D**), where the K-ε-GG motif position indicates the poly-ubiquitin chain type. K48 chains were the most prominent, followed by K63 and K11 chains making up 9.7% and 9.1% of poly-ubiquitin chains in skin respectively. M1 chains were not identified by K-ε-GG proteomics as this chain type is not lysine linked and therefore not enriched by targeting the K-ε-GG motif.

To understand the implications of co-ubiquitinated substrates detected in healthy skin, we performed gene set enrichment analysis of the proteins to which the K-ε-GG peptides belonged. First, we interrogated the cellular components associated with ubiquitinated proteins in healthy skin and observed enrichment of skin specific compartments including melanosome and cornified envelope (**Figure 2E**). Next we used the REVIGO webtool^32^ to reduce the enriched biological processes of ubiquitinated proteins in human skin and displayed them alongside the term dispensability (similarity to other terms) to highlight biological processes that are likely to be mediated by ubiquitination (**Figure 2F**). We found intermediate filament terms, whose proteasomal degradation is known to be regulated by ubiquitination^33,34^, were highly enriched and indispensable confirming this process is widespread and governed by ubiquitination in skin. Novel notable processes enriched in healthy skin were matrix secretion and peptide cross-linking.

An unbiased profile of the skin ubiquitinome gave us the opportunity to predict the upstream enzymes involved in conjugating and removing Ub from target proteins. We utilised the Ubibrowser database^35^, an encyclopaedia of Ub-protein interactions, and by matching the K-ε-GG peptides^36^ detected to the known and predicted E3s and DUBs we generated a prediction of which enzymes are important to Ub dependent processes in healthy skin (**Figure 2G**). We predicted that the most frequent E3 ligases in skin were NEDD4, PRKN, NEDD4L, and SMURF1 all of which have been shown to have Ub dependent roles in regulating cellular processes in skin^37–39^. We predicted that the most frequent DUBs in skin were USP7, USP4, USP8 and CYLD. Taken together, these analyses demonstrate that ubiquitination is integrated into a range of skin processes and likely facilitated by a wide set of E3s and DUBs.

### Increased ubiquitination of CYLD substrates in DUB perturbed CCS tumours

Next, we sought to understand how CYLD perturbation in the skin may disrupt cellular processes. We studied CCS skin tumours, which have biallelic PVs in *CYLD*, preventing its normal DUB activity. We compared the K-ε-GG immunoprecipitation LC-MS/MS data from healthy skin (n=5) to that of CCS skin tumour tissue (n=5). CCS tumours had a similar number of Ub sites; 1508 compared to 1465 K-ε-GG peptides in healthy skin, over 731 proteins but increased ubiquitination levels (**Figure 3A**). We found that the increased ubiquitination could be ascribed to increased mono-ubiquitination with levels of poly-ubiquitin similar between CCS tumours and healthy skin. Additionally, the proportion of poly-ubiquitin chain types present in CCS tumours was not significantly altered from the poly-ubiquitin profile found in healthy skin. Despite this, principal component analysis revealed clustering of the two conditions indicating a distinct difference in their ubiquitinomes and 251 significant DUS (differentially ubiquitinated sites) (t-test, adj.p < 0.05, log2 fold change > 1) (**Figure 3B**). CCS tumours had 151 proteins with upregulated DUS over 217 peptides, including histones, where MACROH2A1 and H1-4 peptides had the greatest increase in ubiquitination. Intriguingly, we also observed increased ubiquitination of skin cell surface receptors including the INSR (9.1 fold change increase compared to normal skin).

**Figure 3.**
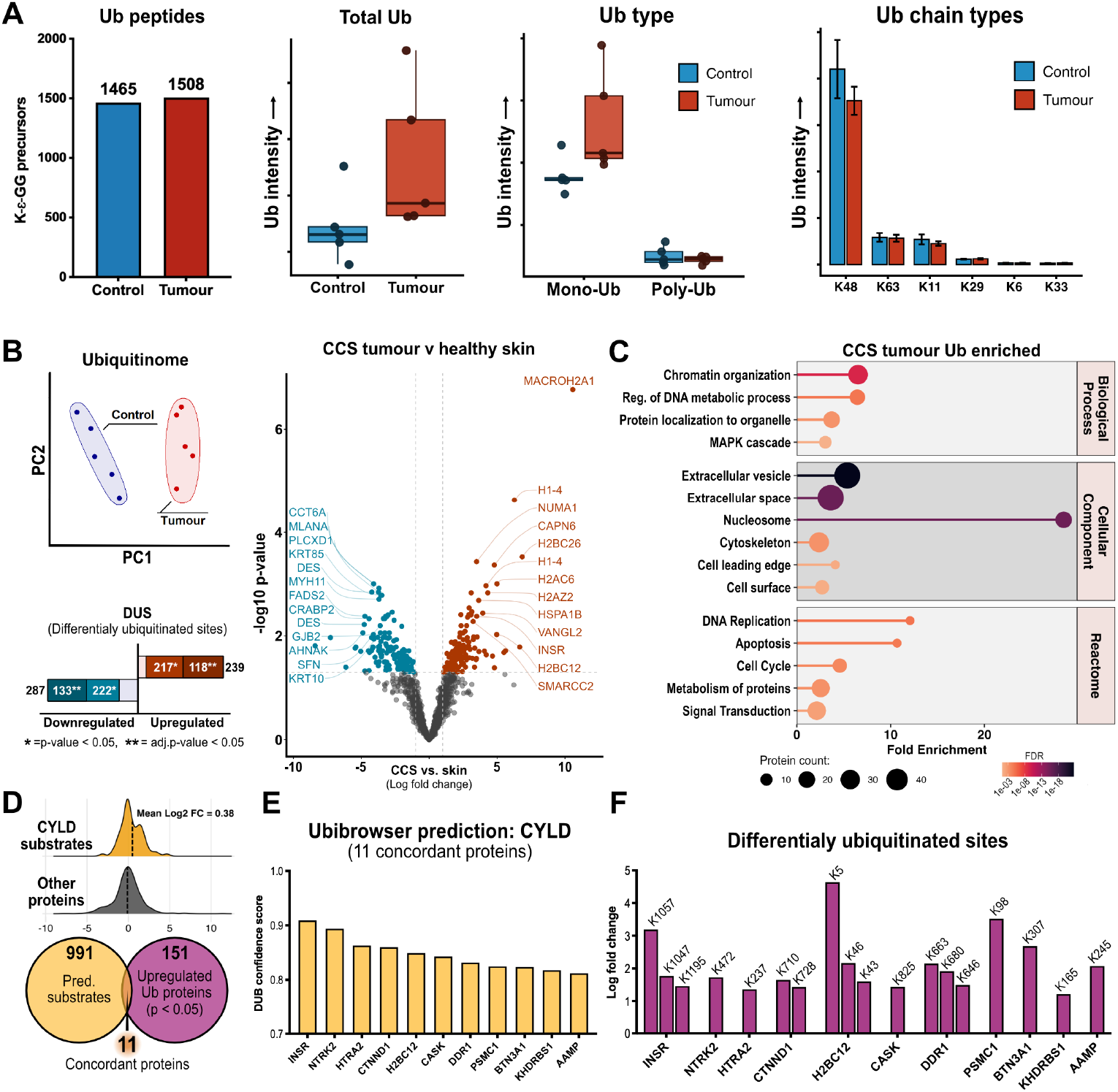
The CCS tumour ubiquitinome. **A)** The ubiquitination status of CCS tumours was compared to control skin. Ubiquitination sites, total ubiquitin, levels of mono- and poly-ubiquitin, and ubiquitin chain types were compared. Box plots display median and upper and lower quartiles. Bar plot error bar display standard error. **B)** Comparison of CCS tumour and control skin K-ε-GG peptide intensities revealed distinct clustering after principal component analysis and differentially ubiquitinated sites (DUS) after t-tests (log fold change >1, p < 0.05 =*, adj.p < 0.05 = **). **C)** Gene set enrichment analysis of DUS in CCS tumours showed significantly altered biological processes, cellular components, and Reactome pathways. Each bar indicates an enriched term, length indicates fold enrichment, colour FDR, and the dot size indicates the number of proteins in the term. **D)** Some ubiquitinated proteins in CCS tumours and healthy skin were also predicted by Ubibrowser to be deubiquitinated by CYLD. The mean log2 fold change of predicted CYLD substrates was compared to the mean log2 fold change of all proteins. **E)** Concordant proteins within the CCS ubiquitinome which had upregulated DUS and were CYLD DUB predicted proteins are displayed alongside **F)** the log2 fold change of their Ub peptides.

We proceeded to use gene set enrichment analysis^40^ to determine the classes of proteins with increased ubiquitination (**Figure 3C**). We identified three main themes of enrichment in the ubiquitinated CCS tumour proteins: (i) Cell division and chromatin organisation, (ii) protein trafficking and extracellular export, and (iii) cellular signal transduction. We also observed increased ubiquitination of secreted vesicles in the extracellular space, and the enriched Reactome pathways highlighted the modulation of mitotic processes, with the pathways M phase, and Mitotic prophase increased in CCS. Additionally, cellular stress response, programmed cell death, and DNA damage pathways were upregulated in the CCS ubiquitinome.

We next wanted to understand the probability that each DUS could be attributed to CYLD perturbation. We interrogated the Ubibrowser database^35^ to identify predicted and known substrates of CYLD. We observed increased ubiquitination in predicted CYLD substrates within the CCS ubiquitinome (mean log2 fold change: 0.38), and within these identified eleven proteins which were significantly upregulated DUS (**Figure 3D**). Of these eleven concordant proteins, the INSR had the highest CYLD substrate prediction score (0.909), followed by other skin relevant proteins NTRK2, CTNND1, and DDR1 (**Figure 3E**). Consistent with this, NTRK2 has been shown to be deregulated in CCS^41^. Further interrogation of DUS within the concordant proteins showed INSR to have three DUS all within the cytoplasmic kinase domain (**Figure 3F**). Furthermore, analysis of predicted INSR DUBs, implicated CYLD as amongst the most likely DUBs, with only USP8 having a higher confidence score (**Supplementary figure 1**).

### Increased INSR ubiquitination in CCS tumours is associated with deregulated basement membrane secretion

Cell surface receptors are central for signal transduction and ubiquitination can affect cellular signalling and receptor recycling. INSR has recognised roles in skin biology including keratinocyte differentiation^42^, wound healing^43^, carcinogenesis^44^, and stimulating hair follicle growth^45^, but has not been studied in CCS tumours. We identified INSR as a DUS in CCS tumours and a predicted CYLD substrate leading us to validate its expression in CCS tumours by immunoblot of tumour lysates and immunostaining of tumour sections (**Figure 4A**). Levels of INSR were increased in CCS tumours compared to skin but varied across patient samples. Spatially, INSR in healthy skin localised to the upper epidermis while in CCS tumours INSR was expressed in central areas of CCS tumour ‘islands’ (dashed line), in patterns that resembled ductal like structures. In both healthy skin and CCS tumours INSR localised to the cell surface, demonstrating its extra-cellular signalling capability.

**Figure 4.**
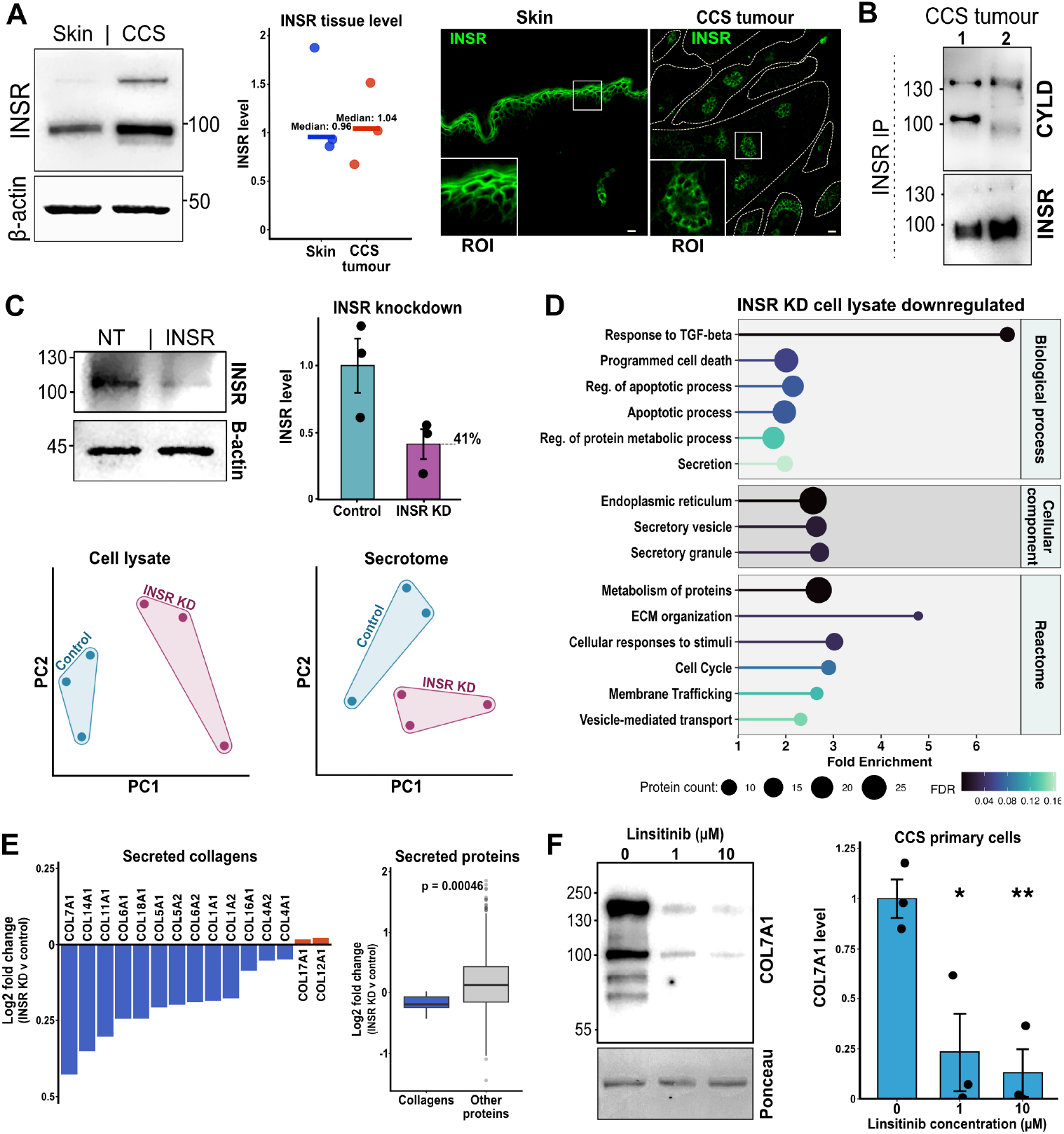
Increased INSR ubiquitination in CCS tumours is associated with basement membrane protein secretion. **A)** Control skin and CCS tumour tissue lysates were probed with anti-INSR antibody by immunoblotting, and tumour sections with anti-INSR, anti-ubiquitin (FK2), and DAPI nuclear stain by immunofluorescence. Region of interest (ROI) displays INSR localisation. Scale bar is 20 μm. **B)** Tissue lysates from CCS tumours were immunoprecipitated with anti-INSR antibody. CYLD co-immunoprecipitation was probed with anti-CYLD (N-terminal) antibody. **C)** CCS tumour keratinocytes were transduced with lentiviral particles targeting INSR (INSR) and with non-targeting particles (NT). Immunoblots against INSR allowed quantification of knockdown (n=3). Error bars represent standard error of the mean. The secretome and cell lysate of INSR knockdown and NT primary CCS tumour keratinocytes were analysed by LC-MS/MS revealing grouping by condition after principal component analysis. **D)** The downregulated gene ontology terms from proteins detected by LC-MS/MS were investigated in INSR knock-down CCS tumour keratinocytes. **E)** Secretomes from CCS tumour keratinocytes were analysed by LC-MS/MS and collagen levels were compared in cells transduced with INSR targeting and non-targeting lentiviral particles. **F)** Primary cultures of CSS tumour keratinocytes were treated with (0,1,10 μM) of INSR inhibitor Linsitinib for 72 h and the culture media collected. Immunoblotting demonstrated a significant reduction in soluble collagen 7 after treatment. Error bars represent standard deviation, p-value <0.05 = *, p-value <0.01 = **.

To demonstrate whether CYLD and INSR interact in CCS tumours, we immunoprecipitated INSR from CCS whole tumour tissue lysates and probed membranes with an N-terminal CYLD antibody (**Figure 4B**). We identified co-immunoprecipitated CYLD at ∼100 kDa in one CCS tumour sample and in another observed evidence of a recently described truncated CYLD protein corresponding to this patient’s pathogenic variant^46^, supporting INSR-CYLD interaction in CCS tumour lysates.

Next, we wanted to determine whether INSR could influence the hypersecretive behaviour of CCS tumours which have been described to produce large amounts of COL7A1^47^. Therefore, we transduced CCS primary tumour keratinocytes with INSR targeting lentiviral particles (INSR KD) and non-targeting (NT) particles (n=3). We reduced INSR levels by over half and studied the resulting secreted and cellular proteins by LC-MS/MS, revealing an altered secretome and proteome after INSR knock-down (**Figure 4C, Supplementary data 3**). Gene set enrichment analysis of the significantly downregulated cellular proteins after INSR knock-down revealed reduced levels of apoptosis regulators and cell cycle proteins, as well as a reduction in proteins associated with extracellular matrix organisation and secretion (**Figure 4D**). This led us to investigate secreted collagen proteins which are hypersecreted in CCS tumours (**Supplementary figure 2**). We observed decreased secretion of collagens after INSR knock-down in CCS tumour keratinocytes, most notably COL7A1 (**Figure 4E**). A significant reduction in collagen proteins in the CCS tumour keratinocyte secretome (one-sample t-test p = 0.00046), coincided with an accumulation of cellular collagens in the cell lysates (**Supplementary figure 3**), suggesting reduced collagen secretion. Further analysis of secreted CCS tumour proteins following INSR knockdown, cross-referenced with the human matrisome database^48^, revealed a selective reduction in collagen secretion (**Supplementary figure 4**).

To further validate the role of INSR in collagen secretion we treated CCS primary tumour keratinocytes with an INSR kinase inhibitor Linsitinib and measured the secretion of COL7A1 in the culture media. We observed a significant reduction in soluble COL7A1 fragments after 1 μM Linsitinib treatment (**Figure 4F**). All detected fragments were reduced, while Ponceau stain and total protein quantification (**Supplementary figure 5**) indicated consistent soluble protein levels suggesting a targeted modulation of COL7A1 secretion. The most prominent fragment (∼170 kDa) corresponds to the previously detected processed alpha chain^49^. Together these findings suggest that a functional CYLD is required for INSR deubiquitination and CYLD PVs in CCS tumours disrupt INSR signalling to promote collagen secretion.

## Discussion

Here we report hundreds of ubiquitinated sites across diverse protein classes and biological processes in healthy human skin, underscoring the importance of Ub in human skin. By utilising CCS tumours as a DUB perturbation model for primarily K63 and M1 ubiquitination in skin^50^, we investigated predicted CYLD substrates and discovered increased INSR ubiquitination at its functional cytoplasmic domain. We sought to modulate INSR function in CCS tumour cells utilising genetic knockdown and chemical inhibition, and found that it was required for CCS tumours basement membrane protein secretion, most notably, COL7A1.

The cutaneous ubiquitinome in healthy skin gave insights into processes involved in the regulation and degradation of multiple substrates responsible for skin homeostasis. We demonstrated using immunofluorescence, enrichment of ubiquitinated substrates at hair follicles and suprabasal keratinocytes, where cells are stratified in differentiation gradients, suggesting a role for regulation of differentiation in skin by ubiquitination. Although many of the roles of ubiquitination may be attributed to K48-associated targeting to the ubiquitin proteasome system, K63 and M1 Ub chains were also identified, which classically regulate signal transduction pathways and protein trafficking^51,52^. Notably, K63 Ub was the second most prevalent Ub linkage in skin, further highlighting its importance to skin homoeostasis. The spatial overlap between K63 ubiquitination in hair follicles and previously reported CYLD expression in the inner root sheath^27^ indicates that CYLD functions to restrict K63 linked ubiquitination in this compartment, providing a mechanistic link between CYLD loss in CCS tumours and hair follicle tumorigenesis.

In CCS tumours, histones had increased ubiquitination including MACROH2A1, H1-4, H2BC6 and H2BC12. MACROH2A1 maintains chromatin in a condensed, transcriptionally repressive state, thereby guiding keratinocyte differentiation programs and cell identity in healthy skin^53^. Ubiquitination and subsequent proteasomal targeting of histones causes chromatin decompaction and increases transcriptional activity which may alter cell identity and promote tumorigenesis. CCS tumours exhibited altered ubiquitination over a range of histones which reflect a global shift in cell state, likely driven by oncogenic changes which promote chromatin decompaction and transcriptional activity originating from mutated tumour suppressor *CYLD*. Enrichment of chromatin organisation, and DNA metabolism biological processes, as well as enrichment of cell cycle related proteins suggests altered regulation of transcription and cell division in CCS tumours.

We observed that CYLD favoured Ub linkages (K63 and M1) were predominantly cytoplasmic in healthy skin, indicating that CYLD activity primarily affects cytoplasmic and membrane bound proteins. We noted increased intracellular receptor ubiquitination of predicted CYLD substrates DDR1, NTRK2, and INSR, which in the case of INSR occurs downstream of INS dependent autophosphorylation^54^. INS and its paralogue insulin-like growth factor 1 (IGF1) are potent stimulators of growth in the skin, regulate hair follicle cycling, and their receptors are influenced by ubiquitination^45,55^. Additionally, CYLD deubiquitinates AKT downstream of INSR^56^, supporting CYLD’s regulation of INSR-AKT signalling at multiple levels.

We identified ubiquitination at K1057 within the INSR kinase active site, a residue that maintains a conserved salt bridge with E1074 to stabilize its active kinase conformation^57^. Studies of the homologue IGF-1R, which shares 100% homology in the activation loop, demonstrate that receptor autophosphorylation is a prerequisite for ubiquitination at this site, which subsequently disrupts the K1057-E1074 salt bridge^57^. Notably, the INSR intracellular domain exhibits ligand and kinase independent signaling that promotes ECM remodelling, cell cycle gene expression, and apoptosis resistance^58^.

We propose that defective CYLD mediated deubiquitination at K1057 prevents insulin dependent receptor inactivation, thereby sustaining INSR signaling. This mechanism could explain the well-established ECM hypersecretory phenotype and apoptosis resistance observed in CCS tumors^16,17^, consistent with our primary cell culture data. Supporting this model, others have shown that cells with INSR/IGF1R double knockout exhibit apoptosis resistance that is reversed upon re-expression of K1057R mutant INSR^59^, indicating that preventing ubiquitination at this site restores apoptotic sensitivity. Furthermore, E1074 mutations that disrupt the K1057-E1074 salt bridge are associated with lung squamous cell neoplasms^60^, indicating an oncogenic effect in disrupting K1057-E1074.

Further work is required to elucidate INSR ubiquitination types, but in the related receptor IGF1R, its K63 ubiquitination by MDM2 promotes receptor internalisation and degradation^61,62^, which may be similar for INSR. These findings lead us to hypothesise that pathogenic *CYLD* variants deregulate INSR signalling enabling increased ECM production.

Our comparative ubiquitinomic analysis establishes ubiquitination as a central regulatory mechanism in human skin and reveals how perturbation of this system drives pathological processes in CYLD deficient tumours. We identify ubiquitination of INSR at K1057 which may contribute to the CCS tumour phenotype, linking aberrant receptor signalling to apoptosis resistance and ECM hypersecretion. Our approach demonstrates the value of ubiquitinomics coupled with database integration, for uncovering dysregulated signalling axis in human tissue, with potential application to other ubiquitin driven disorders.

## Methods

### Primary cell culture

Surplus CCS tumour tissue was cut finely with scalpels and digested with trypsin (40 min, 37 °C) and collagenase 1 mg/mL (overnight, 4 °C). The suspensions were cultured on collagen coated tissue culture plastics in Defined keratinocyte serum free medium (DKSFM, Thermo Fisher Scientific) at 37 °C, 5 % CO2 in a humidified environment.

### Protein extraction

70 μm tissue slices were added to CK mix Precellys tubes (Bertin technologies) with 500 μL of 8M urea lysis buffer. Samples were homogenised at 6800 rpm for 90 s in a Precellys Evolution homogeniser (Bertin technologies). Lysates were transferred to new tubes and sonicated on ice, then centrifuged at 16,000 g for 10 min and soluble lysates collected. Protein quantification was performed by Pierce BCA protein assay (Thermo Fisher Scientific) and measured at 562 nm with SpectraMax iD3 microplate reader (Molecular Devices).

### Peptide generation and di-glycine (K-ε-GG) immunoprecipitation

One milligram of each tissue lysate was reduced with 20 mM TCEP (Tris(2-carboxyethyl)phosphine) for 30 min at 55 °C and alkylated with 20 mM N-ethylmaleimide at RT for 30 min. Samples were diluted to a final concentration of 2 M urea and 20 mM HEPES (4-(2-hydroxyethyl)piperazine-1-ethane-sulfonic acid). Overnight digestion with trypsin was performed at RT with 60 μg of trypsin (Worthington Biochemical). The resulting peptide mix was acidified with formic acid (FA) and desalted on SepPak C18 columns (Waters). Samples were processed with PTMScan HS Ubiquitin/SUMO Remnant Motif (K-ε-GG) kit (Cell Signalling Technology, #59322). The resulting K-ε-GG peptides were dried down and resuspended in 0.1% FA prior to LC-MS/MS analysis.

### LC-MS/MS analysis

Peptides were analysed on a Bruker timsTOF HT MS coupled to a nanoElute II HPLC system (Bruker), operated with an Aurora Elite 15 column (15 cm x 75 μm, 1.7 μm C18, 120 Å pore size, ionoptics) at 23 °C and a 60-minute linear gradient (0 - 35% 100% MeCN, 300 nL/min flowrate). The timsTOF HT was equipped with a CaptiveSpray source (1600 V capillary voltage, 3.0 L/min dry gas, 180 °C dry temperature), and a PepMap100 trap (5 x 0.3 mm, 5 μm C18, Thermo Fisher Scientific). A custom pyDIA-PASEF method was used with 16 variable width windows (ion mobility by *m/z*) with two quadrupole positions per window (*m/z* 420 – 1255, 0.75 - 1.37 1/K_0_, cycle time 2.92 s). The following general TIMS (0.75 - 1.37 1/K_0_, ramp and accumulation times 166 ms each) and MS (m/z 100 – 1700, positive polarity) settings were used and high sensitivity detection was enabled.

### Lentiviral knockdown

To achieve protein knockdown in CCS primary keratinocytes, cultures were transduced with non-targeting or INSR targeting (TCAATGAGGCCGAGGTTGG, TAGACACTGAGATTGGATC, GCGGTACCCGGACAGATGT) shRNA, alongside puromycin resistance and GFP selection marker carrying lentiviral particles for 72 h. GFP positive cells were FACS sorted and recultured.

### Secretome and primary cell proteome analysis

Serum free cell media from primary cell cultures was collected and centrifuged at 16,000 g for 5 min, and the soluble fraction was precipitated with four volumes of −20 °C acetone. Precipitated proteins and primary cell pellets were resuspended in 5% sodium dodecyl sulfate, 50, mM triethylammonium bicarbonate (TEAB) in water, and tryptic peptides were prepared using S-trap micro columns (Protifi) following the manufacturers protocol.

Extended methods section can be found in **Supplementary data 4**.

## Acknowledgements

This work was partly funded by the British Skin Foundation and the Newcastle NIHR Biomedical Research Centre. MG-H is supported by the LEO
Foundation grant number LF18500.

## Data availability

Mass spectrometry proteomics data files are available at the Proteomics Identification database (PRIDE) under the accession identifiers: PXD069159, PXD071326

## Supplementary data and tables

Supplementary data 1 – Healthy skin and CCS tumour ubiquitinome

Supplementary data 2 – Matched reference skin proteome

Supplementary data 3 - Secretome and proteome after INSR knockdown

Supplementary data 4 – Extended material and methods

Supplementary table 1 – Patient samples used in study

Supplementary table 2 – Antibodies used in study

## Author contributions

JI performed all experimental work, data analysis and visualisations, and contributed to experimental design and manuscript writing. AMF operated mass spectrometry instruments. IR contributed to experimental design, data review, and manuscript review and editing. MGH contributed to data review and manuscript review. MT contributed to manuscript review and provided mass spectrometry instrumentation. NR contributed to experimental design, data review, manuscript writing and review.

## References

1. Tracz, M. & Bialek, W. Beyond K48 and K63: non-canonical protein ubiquitination. Cell. Mol. Biol. Lett. 26, 1 (2021).

2. Pinder, J. B., Attwood, K. M. & Dellaire, G. Reading, writing, and repair: the role of ubiquitin and the ubiquitin-like proteins in DNA damage signaling and repair. Front. Genet. 4, 45 (2013).

3. Chen, Z. J. Ubiquitin signalling in the NF-kappaB pathway. Nat. Cell Biol. 7, 758–765 (2005).

4. Dang, F., Nie, L. & Wei, W. Ubiquitin signaling in cell cycle control and tumorigenesis. Cell Death Differ. 28, 427–438 (2021).

5. Erpapazoglou, Z., Walker, O. & Haguenauer-Tsapis, R. Versatile Roles of K63-Linked Ubiquitin Chains in Trafficking. Cells 3, 1027–1088 (2014).

6. Cao, J. & Yan, Q. Histone Ubiquitination and Deubiquitination in Transcription, DNA Damage Response, and Cancer. Front. Oncol. 2, 26 (2012).

7. Oliveira, S. A. et al. Parkin mutations and susceptibility alleles in late-onset Parkinson’s disease. Ann. Neurol. 53, 624–629 (2003).

8. Barker, D. F. et al. BRCA1 R841W: a strong candidate for a common mutation with moderate phenotype. Genet. Epidemiol. 13, 595–604 (1996).

9. Liu, C. et al. Novel Mutations in X-Linked, USP26-Induced Asthenoteratozoospermia and Male Infertility. Cells 10, 1594 (2021).

10. Lessel, D. et al. Dysfunction of the MDM2/p53 axis is linked to premature aging. J. Clin. Invest. 127, 3598–3608 (2017).

11. Lin, Z. et al. Stabilizing mutations of KLHL24 ubiquitin ligase cause loss of keratin 14 and human skin fragility. Nat. Genet. 48, 1508–1516 (2016).

12. Wiesner, T. et al. Germline mutations in BAP1 predispose to melanocytic tumors. Nat. Genet. 43, 1018–1021 (2011).

13. Soysouvanh, F. et al. An Update on the Role of Ubiquitination in Melanoma Development and Therapies. J. Clin. Med. 10, 1133 (2021).

14. Marín-Rubio, J. L., Raote, I., Inns, J., Dobson-Stone, C. & Rajan, N. CYLD in health and disease. Dis. Model. Mech. 16, dmm050093 (2023).

15. Davies, H. R. et al. Epigenetic modifiers DNMT3A and BCOR are recurrently mutated in CYLD cutaneous syndrome. Nat. Commun. 10, 4717 (2019).

16. Tunggal, L. et al. Defective Laminin 5 Processing in Cylindroma Cells. Am. J. Pathol. 160, 459–468 (2002).

17. Bruckner-Tuderman, L., Pfaltz, M. & Schnyder, U. W. Cylindroma overexpresses collagen VII, the major anchoring fibril protein. J. Invest. Dermatol. 96, 729–734 (1991).

18. Tunggal, L. et al. Defective Laminin 5 Processing in Cylindroma Cells. Am. J. Pathol. 160, 459–468 (2002).

19. Sun, S.-C. CYLD: a tumor suppressor deubiquitinase regulating NF-κB activation and diverse biological processes. Cell Death Differ. 17, 25–34 (2010).

20. Moquin, D. M., McQuade, T. & Chan, F. K.-M. CYLD deubiquitinates RIP1 in the TNFα-induced necrosome to facilitate kinase activation and programmed necrosis. PloS One 8, e76841 (2013).

21. Brummelkamp, T. R., Nijman, S. M. B., Dirac, A. M. G. & Bernards, R. Loss of the cylindromatosis tumour suppressor inhibits apoptosis by activating NF-kappaB. Nature 424, 797–801 (2003).

22. Kim, W. et al. Systematic and Quantitative Assessment of the Ubiquitin-Modified Proteome. Mol. Cell 44, 325–340 (2011).

23. Sauerland, M. B. et al. Scalable Acid-Aided Lysis of Skin Samples Improves Proteome Coverage. J. Invest. Dermatol. 0, (2025).

24. Huntzicker, E. G. & Oro, A. E. Controlling hair follicle signaling pathways through polyubiquitination. J. Invest. Dermatol. 128, 1081–1087 (2008).

25. Li, Y. et al. Regulation of epidermal differentiation through KDF1-mediated deubiquitination of IKKα. EMBO Rep. 21, e48566 (2020).

26. Technical report: Targeted proteomic analysis reveals enrichment of atypical ubiquitin chains in contractile murine tissues. J. Proteomics 229, 103963 (2020).

27. Massoumi, R., Podda, M., Fässler, R. & Paus, R. Cylindroma as tumor of hair follicle origin. J. Invest. Dermatol. 126, 1182–1184 (2006).

28. Huntzicker, E. G. & Oro, A. E. Controlling hair follicle signaling pathways through polyubiquitination. J. Invest. Dermatol. 128, 1081–1087 (2008).

29. Hooper, C., Puttamadappa, S. S., Loring, Z., Shekhtman, A. & Bakowska, J. C. Spartin activates atrophin-1-interacting protein 4 (AIP4) E3 ubiquitin ligase and promotes ubiquitination of adipophilin on lipid droplets. BMC Biol. 8, 72 (2010).

30. Bakowska, J. C., Jupille, H., Fatheddin, P., Puertollano, R. & Blackstone, C. Troyer Syndrome Protein Spartin Is Mono-Ubiquitinated and Functions in EGF Receptor Trafficking. Mol. Biol. Cell 18, 1683–1692 (2007).

31. Wojcik, F. et al. Functional crosstalk between histone H2B ubiquitylation and H2A modifications and variants. Nat. Commun. 9, 1394 (2018).

32. Supek, F., Bošnjak, M., Škunca, N. & Šmuc, T. REVIGO summarizes and visualizes long lists of gene ontology terms. PloS One 6, e21800 (2011).

33. Jaitovich, A. et al. Ubiquitin-proteasome-mediated degradation of keratin intermediate filaments in mechanically stimulated A549 cells. J. Biol. Chem. 283, 25348–25355 (2008).

34. Rogel, M. R., Jaitovich, A. & Ridge, K. M. The role of the ubiquitin proteasome pathway in keratin intermediate filament protein degradation. Proc. Am. Thorac. Soc. 7, 71–76 (2010).

35. Wang, X. et al. UbiBrowser 2.0: a comprehensive resource for proteome-wide known and predicted ubiquitin ligase/deubiquitinase-substrate interactions in eukaryotic species. Nucleic Acids Res. 50, D719–D728 (2022).

36. Udeshi, N. D. et al. Refined preparation and use of anti-diglycine remnant (K-ε-GG) antibody enables routine quantification of 10,000s of ubiquitination sites in single proteomics experiments. Mol. Cell. Proteomics MCP 12, 825–831 (2013).

37. Sun, L., Amraei, R. & Rahimi, N. NEDD4 regulates ubiquitination and stability of the cell adhesion molecule IGPR-1 via lysosomal pathway. J. Biomed. Sci. 28, 35 (2021).

38. González-Casacuberta, I. et al. Transcriptional alterations in skin fibroblasts from Parkinson’s disease patients with parkin mutations. Neurobiol. Aging 65, 206–216 (2018).

39. Liu, H. et al. E3 Ubiquitin Ligase NEDD4L Negatively Regulates Skin Tumorigenesis by Inhibiting IL-6/GP130 Signaling Pathway. J. Invest. Dermatol. 144, 2453-2464.e11 (2024).

40. Ge, S. X., Jung, D. & Yao, R. ShinyGO: a graphical gene-set enrichment tool for animals and plants. Bioinformatics 36, 2628–2629 (2020).

41. Rajan, N. et al. Dysregulated TRK signalling is a therapeutic target in CYLD defective tumours. Oncogene 30, 4243–4260 (2011).

42. Differential Roles of Insulin Receptor and Insulin-Like Growth Factor-1 Receptor in Differentiation of Murine Skin Keratinocytes. J. Invest. Dermatol. 115, 24–29 (2000).

43. Liu, Y., Petreaca, M., Yao, M. & Martins-Green, M. Cell and molecular mechanisms of keratinocyte function stimulated by insulin during wound healing. BMC Cell Biol. 10, 1 (2009).

44. Weingarten, G. et al. Insulin receptor plays a central role in skin carcinogenesis by regulating cytoskeleton assembly. FASEB J. 33, 2241–2251 (2019).

45. Philpott, M. P., Sanders, D. A. & Kealey, T. Effects of insulin and insulin-like growth factors on cultured human hair follicles: IGF-I at physiologic concentrations is an important regulator of hair follicle growth in vitro. J. Invest. Dermatol. 102, 857–861 (1994).

46. Hodgson, K. et al. Targeting non-canonical NF-κB signalling in CYLD cutaneous syndrome by selective inhibition of IκB kinase alpha. 2025.01.31.635629 Preprint at 10.1101/2025.01.31.635629 (2025).

47. Bruckner-Tuderman, L., Pfaltz, M. & Schnyder, U. W. Cylindroma overexpresses collagen VII, the major anchoring fibril protein. J. Invest. Dermatol. 96, 729–734 (1991).

48. Naba, A. et al. The Matrisome: In Silico Definition and In Vivo Characterization by Proteomics of Normal and Tumor Extracellular Matrices *. Mol. Cell. Proteomics 11, (2012).

49. Bentz, H. et al. Isolation and partial characterization of a new human collagen with an extended triple-helical structural domain. Proc. Natl. Acad. Sci. U. S. A. 80, 3168–3172 (1983).

50. Komander, D. et al. The structure of the CYLD USP domain explains its specificity for Lys63-linked polyubiquitin and reveals a B box module. Mol. Cell 29, 451–464 (2008).

51. Lim, K. L. et al. Parkin Mediates Nonclassical, Proteasomal-Independent Ubiquitination of Synphilin-1: Implications for Lewy Body Formation. J. Neurosci. 25, 2002–2009 (2005).

52. Emmerich, C. H. et al. Activation of the canonical IKK complex by K63/M1-linked hybrid ubiquitin chains. Proc. Natl. Acad. Sci. 110, 15247–15252 (2013).

53. Creppe, C. et al. MacroH2A1 regulates the balance between self-renewal and differentiation commitment in embryonic and adult stem cells. Mol. Cell. Biol. 32, 1442–1452 (2012).

54. Ahmed, Z., Smith, B. j & Pillay, T. s. The APS adapter protein couples the insulin receptor to the phosphorylation of c-Cbl and facilitates ligand-stimulated ubiquitination of the insulin receptor. FEBS Lett. 475, 31–34 (2000).

55. Sehat, B., Andersson, S., Vasilcanu, R., Girnita, L. & Larsson, O. Role of Ubiquitination in IGF-1 Receptor Signaling and Degradation. PLoS ONE 2, e340 (2007).

56. Lim, J. H. et al. CYLD negatively regulates transforming growth factor-β-signalling via deubiquitinating Akt. Nat. Commun. 3, 771 (2012).

57. Hubbard, S. R. The Insulin Receptor: Both a Prototypical and Atypical Receptor Tyrosine Kinase. Cold Spring Harb. Perspect. Biol. 5, a008946 (2013).

58. Nagao, H. et al. Unique ligand and kinase-independent roles of the insulin receptor in regulation of cell cycle, senescence and apoptosis. Nat. Commun. 14, 57 (2023).

59. Boucher, J. et al. A Kinase-Independent Role for Unoccupied Insulin and IGF-1 Receptors in the Control of Apoptosis. Sci. Signal. 3, ra87–ra87 (2010).

60. Hammerman, P. S. et al. Comprehensive genomic characterization of squamous cell lung cancers. Nature 489, 519–525 (2012).

61. Girnita, L., Girnita, A. & Larsson, O. Mdm2-dependent ubiquitination and degradation of the insulin-like growth factor 1 receptor. Proc. Natl. Acad. Sci. U. S. A. 100, 8247–8252 (2003).

62. Sehat, B., Andersson, S., Girnita, L. & Larsson, O. Identification of c-Cbl as a new ligase for insulin-like growth factor-I receptor with distinct roles from Mdm2 in receptor ubiquitination and endocytosis. Cancer Res. 68, 5669–5677 (2008).

