## Supplementary figure for "Comparative ubiquitinomics of human skin reveals insulin receptor ubiquitination as a regulator of collagen secretion"

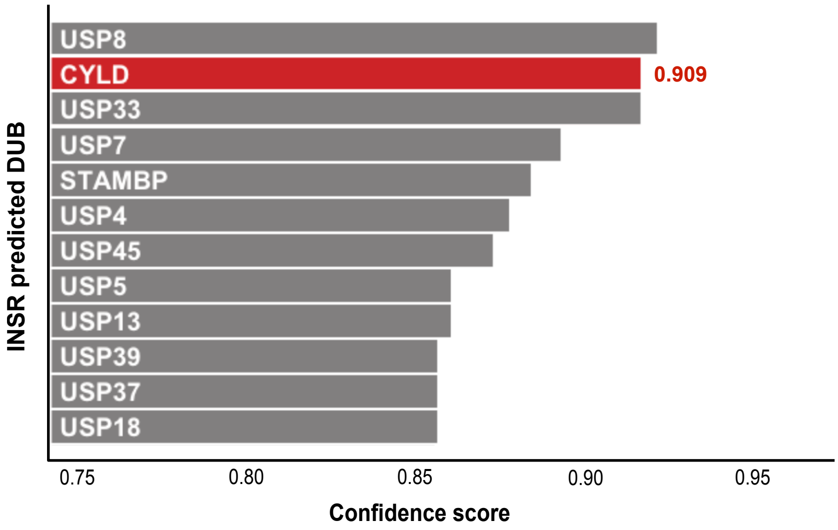


**Supplementary figure 1. CYLD is predicted to deubiquitinate INSR.**
CCS primary cells were cultured in defined keratinocyte serum free media and the protein levels were calculated by bicinchoninic acid (BCA) assay after Linsitinib treatment.

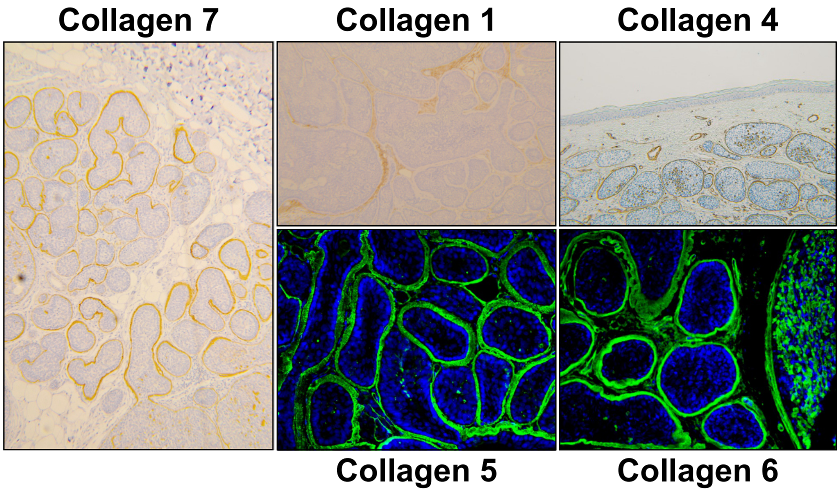


**Supplementary figure 2. Collagen proteins are hypersecreted in CCS tumours.** Immunohistochemistry and immunofluorescence imaging of collagen 1, 4, 5 ,6 and 7 in CCS tumour tissue.

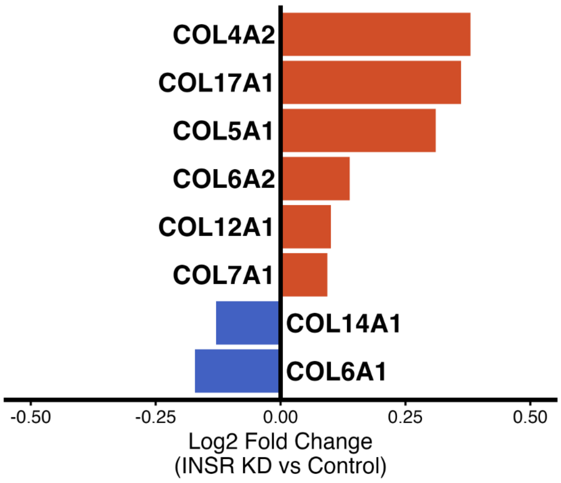


**Supplementary figure 3. Collagen accumulates in CCS tumour keratinocytes after INSR knockdown.** CCS primary keratinocytes were transduced with *INSR* mRNA targeting lentiviral particles. Cell lysates were analysed by LC-MS and collagen levels were compared to cells transduced with non-targeting lentiviral particles.

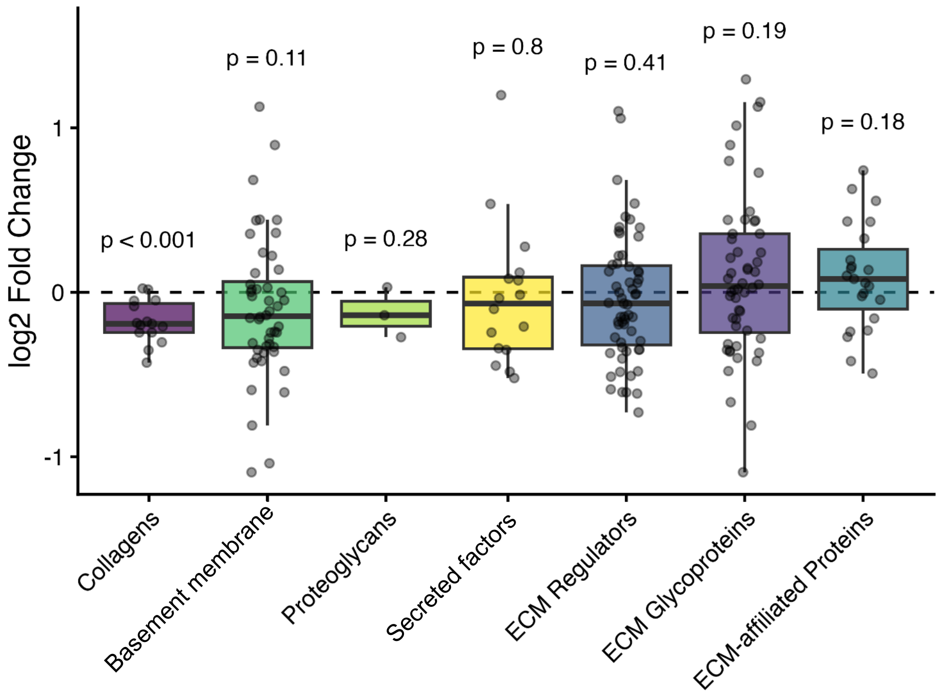


**Supplementary figure 4. Collagens are selectively reduced in the CCS tumour secretome after INSR knockdown.** Analysis of CCS primary cells after INSR knockdown compared to non-targeting control revealed targeted reduction in collagens amongst the human matrisome protein classes. *** = P-value < 0.001 after comparison of the mean to zero using one-sample t-test. ns = non-significant

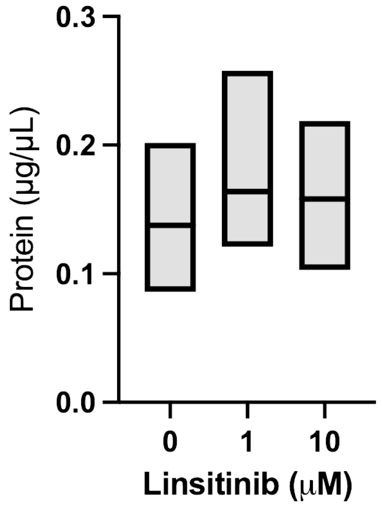


**Supplementary figure 5. Total soluble protein levels in CCS primary cell media.**
CCS primary cells were cultured in defined keratinocyte serum free media and the secreted protein levels were calculated by bicinchoninic acid (BCA) assay after Linsitinib treatment. The median and upper and lower quartiles are shown.

**Supplementary table 1**

Patient samples used in study

| External ID | Age | Sex | CYLD Genotype | Experiment | Comment |
|---|---|---|---|---|---|
| Control 1 | 78 | M | WT | Di-glycine enrichment | Control skin |
| Control 2 | 78 | M | WT | Di-glycine enrichment | Control skin |
| Control 3 | 56 | M | WT | Di-glycine enrichment | Control skin |
| Control 4 | 76 | M | WT | Di-glycine enrichment | Control skin |
| Control 5 | 83 | M | WT | Di-glycine enrichment | Control skin |
| CCS 1 | 44 | F | c.2460delC | Di-glycine enrichment | CCS tumour |
| CCS 2 | 44 | F | c.2460delC | Di-glycine enrichment | CCS tumour |
| CCS 3 | 63 | F | c.2299A>T | Di-glycine enrichment | CCS tumour |
| CCS 4 | 63 | F | c.2299A>T | Di-glycine enrichment | CCS tumour |
| CCS 5 | 44 | F | c.2460delC | Di-glycine enrichment | CCS tumour |
| CCS 6 | 71 | F | c.2460delC | INSR immunopurification | CCS tumour |
| CCS 7 | 49 | F | c.2460delC | INSR immunopurification | CCS tumour |
| CCS 8 | 61 | F | c.2460delC | Linsitinib treatment | CCS primary cells |
| CCS 9 | 61 | F | c.2460delC | Linsitinib treatment | CCS primary cells |
| CCS 10 | 75 | F | c.2441dupA | Linsitinib treatment | CCS primary cells |

**Supplementary table 2.**
Antibodies used in study

| Antibody name | Target | Supplier | Identifier |
|---|---|---|---|
| Anti ubiquitin FK2 | monoubiquitinylated, K29-, K48-, and K63-linked polyubiquitinylated proteins | Merck | ST1200 |
| Anti ubiquitin K63 | K63-linkage specific polyubiquitin | Cell signaling technology | D7A11 |
| Anti ubiquitin M1 | Linear (M1) ubiquitin chains | Merck | MABS451 |
| Anti insulin receptor beta | Residues surrounding Tyr999 of human insulin receptor | Cell signaling technology | 4B8 |
| Anti collagen 7 | N-terminus region of human COL7A1 protein | Cell signaling technology | F4Q8A |
| Anti beta-actin | Beta-actin | Cell signaling technology | D6A8 |
| Anti CYLD | amino acids 1-243 of Human CYLD | Abcam | ab153698 |

**Supplementary data 4. Extended materials and methods**

**Ethics**

Skin samples were provided with signed and informed consent (**Supplementary table 1**). This study was performed under research ethics committee approval from the Hartlepool research ethics committee and north east – Newcastle & north Tyneside 1 research ethics committee (REC Ref: 06/Q1001/59; 08/H0906/95+5; 19/NE/0004).

**Immunofluorescence**

30 µm tissue sections were collected from snap frozen CCS tumours and healthy skin. The sections were fixed on glass slides in ice cold methanol for 10 min, followed by 3 x 10 min washes in PBS. Fixed sections were incubated in 30% sucrose in PBS w/v for 30 min followed by blocking in 0.5% BSA, 0.3% Triton X-100 w/v in PBS for 1 h, RT. Incubation with primary antibody (**Supplementary table 2**) was performed overnight at 4°C. Subsequently, sections were washed 3 x 10 min in 0.2% BSA, 0.1% Triton X-100 w/v in PBS. Fluorescent staining was achieved with AlexaFluor 594 (Thermo Fisher Scientific) conjugated secondary antibody staining for 1 h at RT in the dark. A further 3 x 10 min washes with 0.2% BSA, 0.1% Triton X-100 w/v in PBS were performed, and coverslips were mounted with ProLong Gold antifade with DAPI (Thermo Fisher Scientific). Fluorescence images were captured with a Zeiss Axioimager Z2 microscope.

**Immunoblotting**

Ten micrograms of protein from each sample were mixed with Laemmli buffer, boiled for 5 min at 95 °C, and separated on 10% sodium dodecyl sulphate polyacrylamide gel electrophoresis (SDS-PAGE) gels. Following electrophoresis, the proteins were transferred to a methanol activated polyvinylidene fluoride (PVDF) membrane (Merck). Membranes were blocked with 2 % BSA in TBS-T buffer for 1 h at room temperature. After incubation with primary antibody (**Supplementary table 2**) in 1% BSA in TBS-T buffer, blots were washed with TBS-T at RT for 5 min three times and then incubated with the appropriate horseradish peroxidase (HRP)-conjugated secondary antibody. The peroxidase activity was developed using Amersham ECL western blotting detection reagent. A LiCOR Odyssey® XF imaging system was used for acquisition of images and FIJI (ImageJ) for band densitometry.

**Bioinformatics and statistical analysis**

Proteomic data analysis was performed by searching MS data against an in-silico spectral library generated from the *Homo sapiens* reference proteome (UniProt: UP000005640, downloaded 9$^{th}$ January 2025) using DIA-NN (2.0.2) software on a Terra ubuntu platform. Ub enrichment experiments used an expanded spectral library which included the mass shift indicating the ubiquitin di-glycine remnant. Analysis of peptide and protein group intensities was performed using the R statistical computing environment, including data filtering, clustering, principal component analysis, dendrogram analysis and statistical tests. Gene ontology analysis utilised freely accessible web-based gene set enrichment tools to indicate gene ontology enrichment and statistical confidence from proteins of interest.
